# How many cubs can a mum nurse? Maternal age and size influence litter size in polar bears

**DOI:** 10.1101/532945

**Authors:** Dorinda Marie Folio, Jon Aars, Olivier Gimenez, Andrew E. Derocher, Øystein Wiig, Sarah Cubaynes

## Abstract

Life history theory predicts that females’ age and size affect the level of maternal investment in current reproduction, balanced against future reproductive effort, maintenance and survival. Using long-term (30 years) individual data on 193 female polar bears (*Ursus maritimus*), we assessed age- and size-specific variation on litter size. Litter size varied with maternal age, younger females had higher chances of losing a cub during their first months of life. Results suggest an improvement of reproductive abilities early in life due to experience with subsequent reproductive senescence. Litter size increased with maternal size, indicating that size may reflect individual quality. We also found an optimum in the probability of having twins, suggesting stabilizing selection on female body size. Heterogeneity was observed among the largest females, suggesting that large size comes at a cost.

## 1. Introduction

Life history theory predicts that an optimal level of parental investment should maximize current reproductive success (RS) balanced against maintenance, survival, and future reproduction [1,2]. Among mammals, capital breeders are characterized by high maternal investment [10]. Lactation imposes high energetics demands on mothers [10] whereas energy is stored before breeding when foraging is constrained during reproduction [2]. Mothers’ traits, namely age and body condition, should influence their ability to provide for their young, therefore influencing RS.

RS should increase with age due to an increase in maternal allocation to reproduction as residual reproductive value decreases (“terminal investment” hypothesis [1]). However, recent studies suggest a decline in RS with old age in wild vertebrates because of fewer resources to allocate to reproduction (“reproductive senescence” hypothesis [5]). An increase in RS has also been observed early in life due to increasing breeding abilities with experience (“constraint hypothesis” [6]). Moreover, irrespective of age, RS can vary with female body size and mass [3,4]. Larger size might benefit reproduction by improving or reflecting foraging abilities [7] and lactation [8]. RS might therefore increase until an optimal maternal size or age and then potentially decrease or level off because maintenance costs exceed the benefits associated with higher size or experience of older females [3,9]. However, to date, the influence of maternal traits on reproductive outputs in mammalian capital breeders has received little attention [3].

Using long-term individual data, we assessed age- and size-specific variation in female polar bears (*Ursus maritimus*) relative to litter sizes, which vary from one to three young, in the Svalbard population. We assumed high maternal investment because polar bears i) live in an extreme environment, ii) rely only on stored fat reserves during pregnancy and for the first four months of lactation, and iii) continue to care for and feed young during two and a half years [11]. We expected litter size to increase until an optimal age and size, due to experience and individual quality, and then decline for the largest, and for senescent, individuals.

## 2. Materials and methods

### (a) Data collection

We live-captured polar bears from 1992 to 2017 at Svalbard, Norway, from late March to beginning of May – just after females have emerged from maternity dens with their cubs [11] – using methods described in Stirling et al. 1989 [12]. The age of first reproduction for Svalbard females is usually six years of age [11]. Age was estimated using a premolar tooth extracted from sub-adult and adult bears (cubs were of known age based on size) [13]. Body straight length (cm), hereafter size, was measured as the dorsal straight-line made from the tip of the nose to the caudal end of the tail bone with bears laying sternally recumbent. Litter size (one to three) was recorded upon capture as the number of cubs-of-the-year a mother had reared to that time.

### (b) Statistical analyses

We analysed litter size as a function of mother’s traits using multinomial regression [14] that extends standard logistic regression to more than two outcomes. Litters of one were chosen as the reference category. We did not consider whole litter loss here, and we included only females that were observed with at least one young in the analyses. Separate odd-ratios (OR) were determined for the relative risk of a litter size of “two” *versus* “one”, and the relative risk of a litter size of “three” *versus* “one”, as a function of the covariates. Parameters α and γ respectively represent the OR of twins and triplets, and will give an estimated probability of having one, two or three cubs as a function of the covariates. We tested for linear and quadratic effects of maternal age and size. Because fieldwork was spread over more than a month and that mortality rates for cubs within their first year can be high [11], we considered capture date (as an ordinal date with 1^st^ January being 1 in normal years and 0 in leap years) to account for a possible loss of young between den emergence and observation. A yearly random effect was included on α and γ to account for environmental variation. Twenty-seven models (Table 1) were fitted with a Bayesian approach using Markov Chain Monte Carlo (MCMC) techniques in JAGS [15]. We used non-informative normal prior distributions for the regression coefficients and a uniform prior distribution for the standard deviation of the random effect. We ran two MCMC in parallel with different initial values, 200,000 iterations each and an initial burn-in of 40,000 iterations. One out of ten values were kept. We assessed convergence by visual inspection and by using the Gelman and Rubin R-hat diagnostic (R-hat < 1.1 [16]). For model comparison, we used the Deviance Information Criteria (DIC [17]) and considered the model with lowest DIC as being best supported by the data. Model selection consisted of 5 steps. In step 1, we compared different model structures for the intercept (same or different intercept on α and γ). In step 2 (respectively 3 and 4) we tested for the effect of maternal age (respectively maternal size and capture date). Each tested variable could influence differently both OR (1: one common coefficient, 2: two distinct ones, 3: one for α or 4: one for γ). In step 5, we compared models with additive effects and interactions between the previously selected variables. For ease of interpretation, we considered young females to be aged between 6 and 9 years old because this should be age at their first reproduction; passed 15 years old, we considered females as being old because previous studies suggested reproductive and body senescence around that age [18]; last, we considered prime-aged females as being aged from 10 to 15 years old. Using these cut-offs, we fitted an additional model including age as a factor in the model best supported by the data to assess differences in each age class (Table 2).

**Table 1:** List of all models considered with comparison based on Deviance Information Criteria (DIC) values, with ΔDIC for the difference between the model with two intercepts only and the model under investigation. Parameters α and γ represent the odds ratio (OR) of twins and triplets, respectively, while # par. is for the number of model parameters. For each step, the model best supported by the data is in bold.

| Steps | Models | $\alpha$ twins OR | $\gamma$ triplets OR | # par. | DIC | ΔDIC |
| --- | --- | --- | --- | --- | --- | --- |
| 1: Intercept | **1.1** | **Intercept** | **Intercept '** | **2** | **365.8** | **0** |
|  | 1.2 | Intercept | Intercept | 1 | 515.7 | 150.5 |
| 2: Age | **2.1** | **Age<sup>2</sup>** | **$\beta_0'$** | **4** | **353.5** | **-12.3** |
|  | 2.2 | Age<sup>2</sup> | Age<sup>2</sup> ' | 6 | 356.5 | -9.3 |
|  | 2.3 | Age<sup>2</sup> | Age<sup>2</sup> | 4 | 359.4 | -6.4 |
| | 2.4 | $\beta_0$ | Age' | 3 | 364.7 | -1.1 |
|  | 2.5 | Age | Age' | 4 | 365.5 | -0.3 |
| | 2.6 | Age | $\beta_0'$ ' | 3 | 365.5 | -0.3 |
| | 2.7 | $\beta_0$ | Age<sup>2</sup> ' | 4 | 366.4 | 0.6 |
|  | 2.8 | Age | Age | 3 | 367 | 1.2 |
| 3: Size | **3.1** | **Size<sup>2</sup>** | **Size'** | **5** | **360.3** | **-5.5** |
| | 3.2 | $\beta_0$ | Size' | 3 | 361.4 | -4.4 |
|  | 3.3 | Size<sup>2</sup> | Size<sup>2</sup> ' | 6 | 362 | -3.8 |
| | 3.4 | $\beta_0$ | Size<sup>2</sup> ' | 4 | 362.1 | -3.7 |
| | 3.5 | Size<sup>2</sup> | $\beta_0'$ | 4 | 362.9 | -2.9 |
|  | 3.6 | Size | Size' | 4 | 363.2 | -2.6 |
| | 3.7 | Size | $\beta_0'$ | 3 | 366.8 | 1 |
|  | 3.8 | Size<sup>2</sup> | Size<sup>2</sup> | 4 | 367.5 | 1.7 |
|  | 3.9 | Size | Size | 3 | 367.9 | 2.1 |
| 4: Date | **4.1** | **Date** | **$\beta_0'$** | **3** | **363.7** | **-2.1** |
|  | 4.2 | Date | Date | 3 | 364.6 | -1.2 |
|  | 4.3 | Date | Date' | 4 | 365.8 | 0 |
| | 4.4 | $\beta_0$ | Date' | 3 | 367.4 | 1.6 |
| 5: Additive effects and interactions | 5.1 | Age<sup>2</sup> + Date + Size<sup>2</sup> | Size' | 8 | 346.9 | 0 |
|  | **5.2** | **Age<sup>2</sup> x Date + Size<sup>2</sup>** | **Size'** | **10** | **342.8** | **-4.1** |
|  | 5.3 | Age<sup>2</sup> + Date * Size<sup>2</sup> | Size' | 10 | 348.6 | 1.7 |
|  | 5.4 | Age<sup>2</sup> x Size<sup>2</sup> + Date | Size' | 12 | 354.1 | 7.2 |

**Table 2:** Parameter estimates of the multinomial regression model derived from the model best supported by the data (see Table 1) using age as a factor. Posterior mean, standard deviation (SD) and 95% credible intervals are provided for the odds ratios (fixed effects) as well as the variance of the random effect year. Rhat is the potential scale reduction factor (at convergence, Rhat < 1.2).

|  | Parameter | Mean | SD | 95% credible interval | Rhat |
| --- | --- | --- | --- | --- | --- |
| Fixed effects |  |  |  |  |  |
| $\alpha$ | Intercept | 1.399 | 0.275 | [0.881; 1.962] | 1.001 |
|  | Age (young) | -0.653 | 0.350 | [-1.352; 0.023] | 1.001 |
|  | Age (old) | -1.183 | 0.421 | [-2.015; -0.356] | 1.001 |
|  | Date | -0.047 | 0.207 | [-0.451; 0.357] | 1.001 |
|  | Size | -0.005 | 0.158 | [-0.312; 0.312] | 1.001 |
|  | Size<sup>2</sup> | -0.279 | 0.112 | [-0.505; -0.062] | 1.001 |
|  | Age (young) x Date | -0.760 | 0.390 | [-1.550; -0.017] | 1.001 |
|  | Age (old) x Date | -0.149 | 0.404 | [-0.946; 0.638] | 1.001 |
| $\gamma$ | Intercept' | -2.634 | 0.486 | [-3.671; -1.772] | 1.001 |
|  | Size' | 0.732 | 0.358 | [0.056; 1.466] | 1.001 |
| Year random effect |  |  |  |  |  |
|  | Variance | 0.398 | 0.246 | [0.024; 0.936] | 1.001 |

## 3. Results

The best model (model 5.2, Table 1) included an interaction between a quadratic effect of age and capture date within the field season on the probability of having a litter of two over one cub.

Maternal size influenced both the probability of having two over one cub, and three over one cub. Concerning age, the probability of having twins increased and then decreased after mid-season (day 105) (Figure 1, ESM Figure 1).

**Figure 1:**
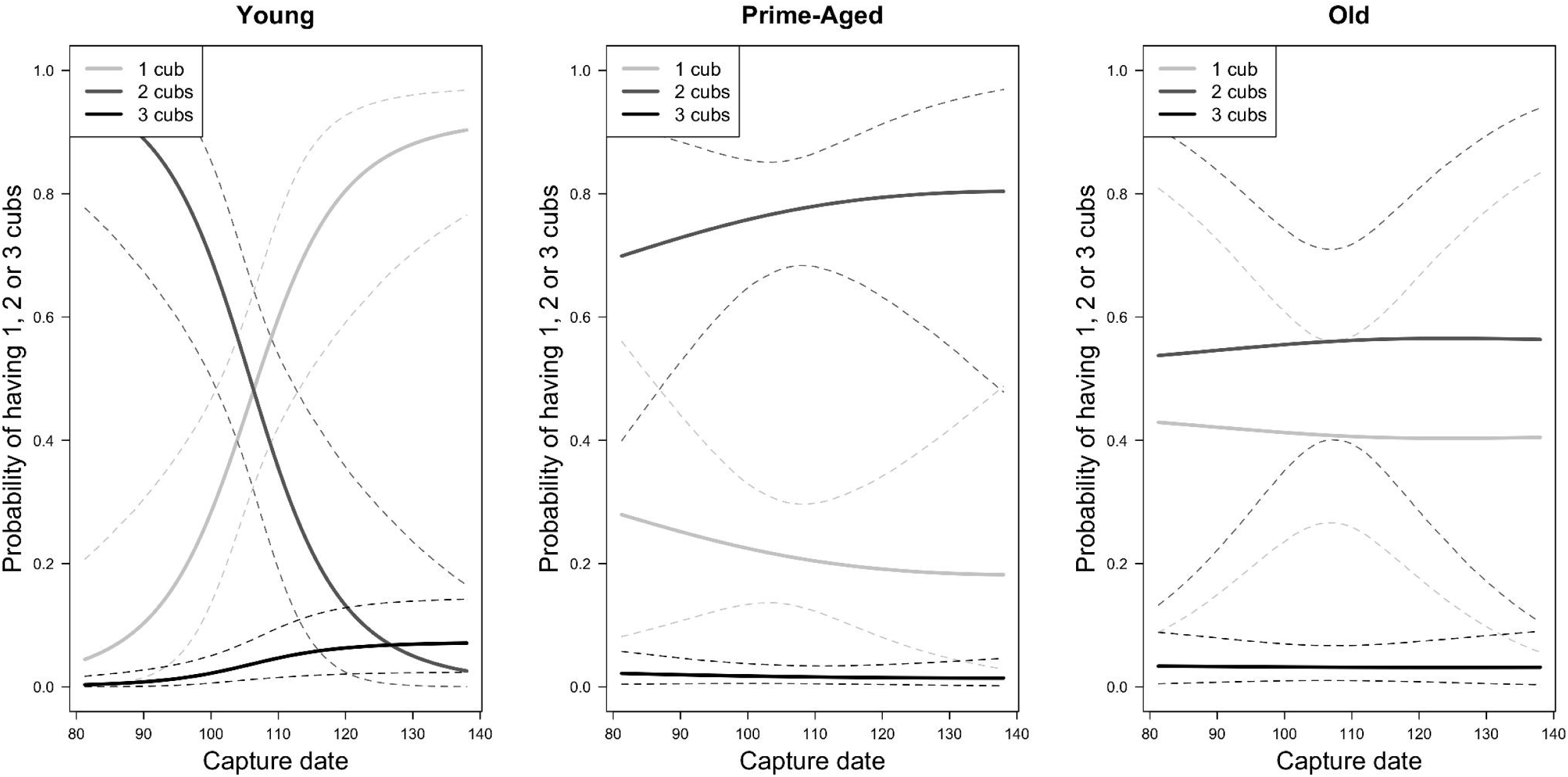
Estimated probability of having 1, 2 or 3 cubs as a function of capture date (in days from March 21^st^) for (a) young (6 years old), (b) prime-aged (12 years old) and (c) old (18 years old) mothers and for a mean maternal size (194.8 cm). Predictions were obtained from the best model (model 5.2 in Table 1). Solid lines are posterior means while dotted lines are 95% credible intervals.

Young females (6-9 years old (y.o.)) had a high probability of having twins shortly after denning (day 90, P(y=2)≈0.9), but it declined within the field season, leading to a higher probability of having just one cub alive towards the end of it (after day 125, P(y=1)≈0.9, Figure 1a). Early in the season, prime-aged females (10-15 y.o.) also had a higher probability of having two over one cub (P(y=2)=0.7 and P(y=1)=0.3, Figure 1b) while older females (>15 y.o.) had similar probabilities of having singletons or twins (P(y=2)≈P(y=1)≈0.5, Figure 1c). In contrast to the marked drop in spring litter size with time for young females, little variation was observed for prime-aged (0.7<P(y=2)<0.8 and 0.2<P(y=1)<0.3, Figure 1b) and old females (P(y=2)≈P(y=1)≈0.5, Figure 1c). Examining parameter estimates confirmed that the interaction term between age and date was only significant for young mothers and not for older ones (Table 2).

Litter size globally increased with the size of a mother (Figure 2). A higher probability of having singletons was found for smaller females (size<183cm, 0.5<P(y=1)<0.8, 0.2<P(y=2)<0.5, P(y=3)=0), of having twins for medium-sized females (190cm<size<200cm, P(y=1)=0.3, P(y=2)=0.7, P(y=3)=0), and of having singletons or triplets for larger females (size>210 cm, P(y=1)≈0.5, 0<P(y=2)<0.2, 0.2<P(y=3)<0.5). However, sample size was very low for triplets (n=8).

**Figure 2:**
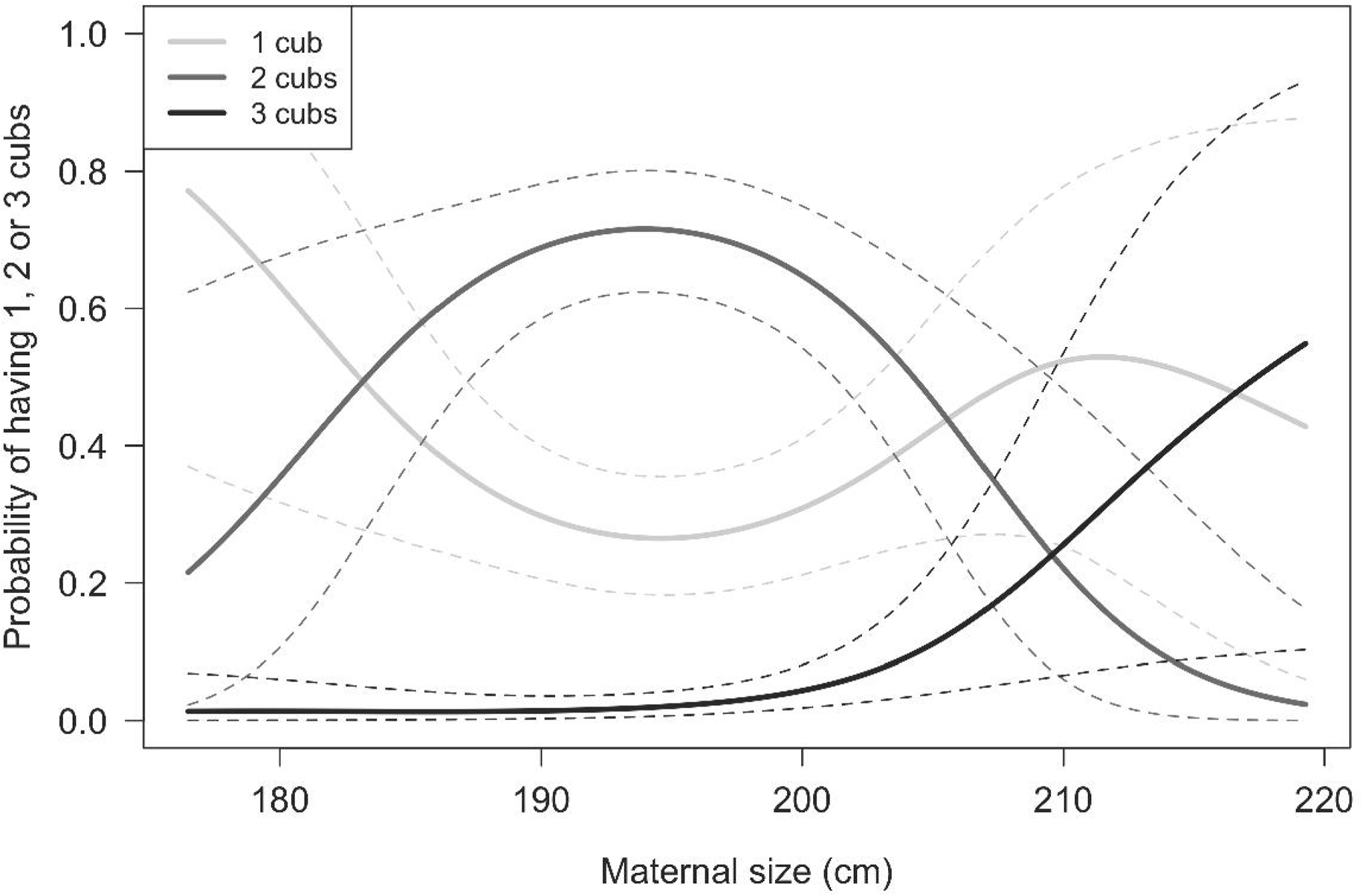
Estimated probability of having 1, 2, or 3 cubs as a function of maternal size for a mean value of mother’s age (11.4 years) and capture date (day 105 ∼ 16^th^ of April). Predictions were obtained from the best model (model 5.2 in Table 1). Solid lines are posterior means while dotted lines are 95% credible intervals.

## 4. Discussion

Our results showed an influence of maternal traits on litter size, an index of reproductive success, suggesting that a mothers’ ability to invest in reproduction and care for their offspring varied with their age and size. The quadratic pattern of variation in litter size observed with age supported the hypothesis of a benefit of gaining experience early in life [6] until 12 y.o. and of reproductive senescence [5] starting from 15 y.o. These results contradict the terminal investment hypothesis [1], and support results on Canadian polar bears showing a decrease in litter size and maternal body senescence after 16 years [18].

Under the experience hypothesis, an improvement of female’s hunting skills might explain the improvement of reproductive investment early in reproductive life [19]. Other studies on mammals [3,18] suggest that an age-related increase in reproductive success could be linked to mass gain and hence resource availability. Further supporting the experience hypothesis, we found that younger females (< 10 y.o.) had about the same probability as prime-aged females to produce two cubs, but higher chances to lose one during the capture spring season.

Under the senescence hypothesis, degradation of physiological functions with aging [20] might impair females’ fat stores accumulation, causing a simultaneous decrease in females’ mass and reproductive outcomes [18]. Older mothers might therefore acquire less energy, and might have higher energy allocation needs toward self-maintenance, reducing energy allocated toward reproduction [1]. Reproductive senescence has been documented in many wild populations for several reproductive parameters such as litter size [4,18], offspring mass [3,5,18], and survival [4].

We showed that, on average, litter size increased with maternal size. The probability of having triplets was only shown to depend on mother’s size, although the sample size for triplets was small. Large size might therefore be an index of individual quality, like in wolves (*Canis lupus*) [4]. The increase in the probability of having twins, and decrease in probability of having singletons, for females up to an optimum size, support this idea. Other traits highlighting foraging capacities have been related to RS – e.g. body mass and condition in bears and other species [3,18]. Among the largest females, the chances of having a singleton or triplets were almost equal and increased, while that of having twins was low and decreasing. Considering the decrease in the probability of having twins, and the increase in the probability of singletons, stabilizing selection on adult female body size is likely to happen: increased investment in growth likely comes at a cost in terms of somatic maintenance.

Considering results on triplets, we suspect higher individual heterogeneity among the largest females compared to smaller and medium sized ones. With a larger sample size, this could be tested by assessing variance in litter size probability. Results on litter size probability suggest that there is one group of ‘good quality females’ having triplets and another group of ‘low quality females’ having singletons. For the latter group, larger size could be associated with a cost that may depend on other factors, such as body condition and environmental quality. Large individuals occupying resource-poor habitats, or experiencing a year of reduced resource availability, might not have enough resources to allocate to both their own maintenance and care for triplets. Heterogeneity in individual quality may override reproductive cost [22], and costs may be restricted to resource-limited contexts [23].

Overall, we found that litter size in polar bears increased with age of mothers early in life until a plateau, followed by a decrease for old females. Because population growth mostly depends on female’s RS, itself influenced by maternal traits, our findings highlight the importance of accounting for individual heterogeneity to understand the species response to environmental perturbations. Future research will aim at understanding the determinants of female polar bears’ reproductive tactics by accounting for environmental conditions. Influence of climate variability has been shown to affect reproductive parameters in several populations, including Svalbard [24].

## Supporting information

ESM1

## Acknowledgements

We thank Magnus Andersen for his participation in fieldwork, and Thor Larsen for initiating the monitoring.

## Authors’ contributions

S.C., J.A. and O.G. conceived the analysis. J.A., AED and ØW collected the data. D.F. carried out the analyses, and S.C., O.G., J.A. advised the analysis. D.F. drafted the manuscript. All authors revised the manuscript critically, gave final approval for publication and agreed to be accountable for all aspects of the work.

## Data accessibility

Data are archived in the Dryad Digital Repository at https://datadryad.org/review?doi=doi:10.5061/dryad.mb2d353.

## Ethical statement

Animal handling methods were approved by the National Animal Research Authority (http://www.mattilsynet.no/fdu).

