## Supplementary material for "How many cubs can a mum nurse? Maternal age and size influence litter size in polar bears": ESM1

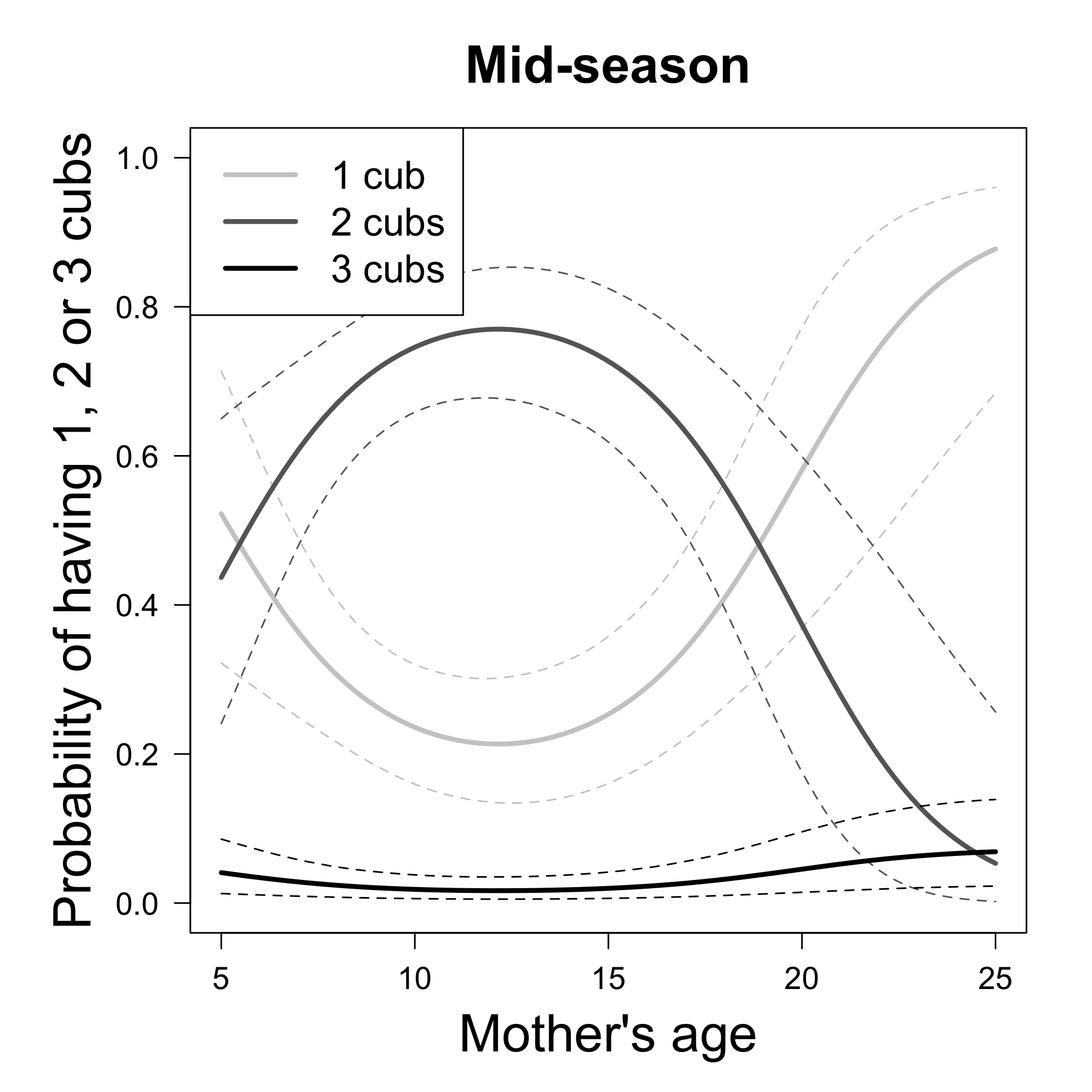

Estimated probability of having 1, 2 or 3 cubs as a function of age (in years) at mid-season (day 105) for a mean maternal size (194.8cm). Predictions were obtained from the best model (model 5.2 in Table 1). Solid lines are posterior means while dotted lines are 95% credible intervals.
